# Deep Learning Based Identification of Tissue of Origin for Carcinomas of Unknown Primary utilizing micro-RNA expression

**DOI:** 10.1101/2024.01.17.576064

**Authors:** Ananya Raghu, Anisha Raghu, Jillian F. Wise

**Affiliations:** Quarry Lane School, San Ramon, CA; Broad Institute of MIT and Harvard; Tufts University, Pre-College Programs, Medford, MA; Salve Regina University, Newport, RI

## Abstract

Carcinoma of Unknown Primary (CUP) is a subset of metastatic cancers in which the primary tissue source, or origin, remains unidentified. CUP accounts for three to five percent of all malignancies [2]. Representing an exceptionally aggressive category of metastatic cancers, the median survival of those diagnosed with CUP is approximately three to six months [1]. The tissue in which a cancer arises plays a key role in our understanding of altered gene expression, altered cellular pathways, and sensitivities to various forms of cell death in cancer cells [3]. Thus, the lack of knowledge on tissue of origin makes it difficult to devise tailored treatments for patients with CUP [4]. Developing clinically implementable methods to identify the tissue of origin of the primary site is crucial in treating CUP patients [4]. In particular, the expression profiles of non-coding RNAs can provide insight into the tissue of origin for CUP. Non-coding RNAs provide a robust route to clinical implementation due to their resistance against chemical degradation [5].

In this work, we investigate the potential of microRNAs as highly accurate biomarkers for detecting the tissue of origin for metastatic cancers. We further hypothesize that data driven approaches can identify specific microRNA biomarker targets. We used microRNA expression data from the Cancer Genome Atlas (TCGA) dataset [6] and assessed various machine learning approaches. Our results show that it is possible to design robust classifiers to detect the tissue of origin for metastatic samples on the TCGA dataset with an accuracy of up to 96%, which may be utilized in situations of CUP. As a validation of our classifiers, we evaluated the accuracy on a separate set of 194 primary tumor samples from the Sequence Read Archive (SRA) [7]. Our findings demonstrate that deep learning techniques enhance prediction accuracy. We progressed from an initial accuracy prediction of 62.5% with decision trees to 93.2% with logistic regression, finally achieving 96.1% accuracy using deep learning on metastatic samples. On the SRA validation set, a lower accuracy of 41.2% was achieved by decision tree, while deep learning achieved a higher accuracy of 81.2%. Notably, our feature importance analysis showed the top three important biomarkers for predicting tissue of origin to be mir-10b, mir-205, and mir-196b, which aligns with previous work [10]. Our findings highlight the potential of using machine learning techniques to devise tests for detecting tissue of origin for CUP. Since microRNAs are carried throughout the body via vesicles secreted from cells, they may serve as key biomarkers for liquid biopsy due to their presence in blood plasma [11]. Our work serves as a foundation towards developing blood-based cancer detection tests based on microRNA presence.

## INTRODUCTION

Carcinoma of unknown origin (CUP) occurs when a patient presents at diagnosis with malignant disease across the body, yet the cancer cells tissue of origin remains unidentifiable. Thus, CUP is a unique subset of metastasized cancer representing an advanced stage in which the cancer has gained the ability to thrive in new tissue sites and has spread from the primary tumor site. In the United States, an estimated 31490 people were diagnosed with cases of cancer of unknown tissue of origin in 2008. This accounts for nearly 3-5% of all cancer cases and given the lack of knowledge on tissue response to current therapeutics the median survival of patients remains only 3-9 months [12]. In many cases of CUP the primary site is never identified, preventing the use of treatment that can be effective for the true tissue of origin [13]. It has been demonstrated that pinpointing the primary site can significantly increase survival rates by enabling for precise and targeted treatment [14].

### Prior Work

Unfortunately, primary tumor identification poses various challenges. Techniques such as serum tumor markers and imaging tests are used to identify the tissue of origin, though only 30% of these tests are successful. Moreover, some positive findings can be misleading [15] and CUP diagnostic workups are often time-consuming, expensive, and unsuccessful [16].

These difficulties have spurred interest in utilizing genetic expression data to identify the tissue of origin. MicroRNAs belong to a class of non-coding regulatory RNAs and have shown great potential for identifying tissue of origin for cancers of unknown primary origin [17].

MicroRNAs are small single stranded RNA molecules that are between 19-25 nucleotides long and are involved in the regulation of gene expression of mRNAs. MicroRNAs hold promise as informative biomarkers for cancer due to their significant involvement in cellular processes such as cell division, apoptosis, proliferation and oncogenesis [18]. Beyond their intracellular role in gene regulation, microRNAs may be carried throughout the body via extracellular vesicles secreted from cells and have been identified in the blood. MicroRNAs used as a biomarker are enhanced further by their remarkable stability in plasma and serum, especially in metastatic cancers [19].

Machine learning has gained attention from the oncology research community and is being applied in many facets including imaging analysis and detection of suitable cancer biomarkers [20]. Deep learning is a subset of machine learning designed to mimic the human brain through the use of artificial neural networks. Generally, deep learning techniques are well suited for discovering and recognizing complex patterns in data that traditional machine learning methods can often miss. In some instances, deep learning techniques have achieved an accuracy of 92% in detecting tissue of origin for metastatic tumors using mRNA data [21].

### Hypotheses

In this study, we set out to explore the possibility of using microRNA from metastatic tissues to determine the primary tissue site. Successful primary site detection from microRNAs will provide a route for cancer detection without requiring samples from the primary tumor site in cases of CUP malignancies. We hypothesize that we would be able to predict the origin of metastatic tumors with high accuracy using microRNA expression data. Moreover, we explore whether deep learning techniques could significantly improve accuracy.

### Datasets

The data for this project was collected from the Cancer Genome Atlas (TCGA) dataset [6] and the SRA [7] from miTED. The TCGA dataset contains samples from 18 different cancer types representing 9648 samples, of which 365 were metastatic, 633 were solid normal, and 8650 were from the primary tumor site. Each sample consisted of microRNA expression data, available as RPM (reads per million mapped reads) as well as metadata including age and gender. The validation SRA cohort consisted of 194 samples from five different cancer types, all of which were from the primary tumor.

### Overview

We developed four machine learning models, a decision tree classifier, random forest, logistic regression and finally a deep learning model. Our deep learning model performed with highest accuracy, achieving an accuracy of 96.1% on detecting tissue of origin for metastatic samples and 81.2% on the validation cohort. Feature importance analysis revealed the top three pivotal biomarkers as mir-10b, mir-196b and mir-205, which confirms prior investigations on microRNAs associated with metastatic cancer [10].

## METHODS

In Figure S2, we outline the data preprocessing pipeline. Our study analyzed published data and did not generate any new sequencing data. TCGA data was obtained from [6]. Data was further filtered by querying the GDC via the APIs specified in [24]. We restricted the tissue type to be one of primary tumor, solid tissue normal, or metastatic (Table S1). We further restricted the data as microRNA transcriptome profiling and picked data corresponding to 18 types of cancer each containing a sufficient amount of samples, obtaining 9,648 files.

To obtain the SRA data, we used the miTED portal and restricted the cancer types to six types of cancer, seen in further detail in Table S1. We obtained 207 samples each containing expression data for 2656 microRNAs. After removing samples with missing features, 194 samples were remaining.

Considering the RPM field, we restricted to microRNAs present in at least 50% of the cell lines using python pandas library [25]. This step reduced the number of features in the TCGA dataset from 1889 to 562. We then picked the common features between the SRA dataset and the TCGA dataset, reducing this number to 497. On both datasets, we normalized the TPM of the selected features per sample to sum to a million. We then transformed the TPM values using the transformation log(TPM + 1) to restrict the range of the input.

We split the TCGA dataset into primary tumor/solid normal samples and metastatic samples. We then further split the primary tumor and solid normal samples into a training and validation set with a 9:1 ratio. For implementation of decision tree, random forest, and logistic regression classifiers the sklearn package was used [26]. We used classification accuracy as the primary metric to evaluate our models using sklearn’s implementation of this function. Deep learning models were created with pytorch [27]. To optimize and train our neural network we utilized Adam optimizer and trained for 50 epochs. Since our objective was classification, we used softmax with cross entropy loss to optimize the model. We used the validation set to determine the hyperparameters of the model. Feature importance was calculated with sklearn’s permutation feature importance function.

We share our implementation at [28]. The implementation script for selecting the specific queried data described above is available in gdc_query.py file in our repository. Code for implementing age and microRNA normalization is available in the file process_data.py. The implementation for model training, performance evaluation, and feature importance is available in the notebook miRNA_model_training_eval.ipynb.

## RESULTS

In order to develop a model to detect tissue of origin, we set the training dataset as the TCGA primary tumor site and solid normal tissue cohorts. We created four machine learning models and assessed them on two datasets. The first dataset consists of metastatic tissues from TCGA and the second dataset, a validation set, consists of primary tumor samples from the SRA.

**Table 1:** Accuracy of developed models on metastatic and SRA test sets. The accuracy for all four models is presented on the TCGA metastatic and SRA cohorts. The decision tree classifier utilized a depth of 14 and the random forest a depth of 19.

| Classifier | Accuracy on metastatic test set | Accuracy on SRA test set |
| --- | --- | --- |
| Decision Tree | 62.5% | 41.2% |
| Random Forest | 94.2% | 74.2% |
| Logistic Regression | 93.2% | 71.6% |
| Deep Learning | 96.1% | 81.2% |

The input features of our models consist of microRNA expression data common to TCGA and SRA data sets. Figure 1 describes the overall architecture of the model which consists of 2 linear layers. The second layer has 18 outputs, corresponding to each cancer type. The cancer type corresponds to the output with the maximum value.

**Figure 1:**
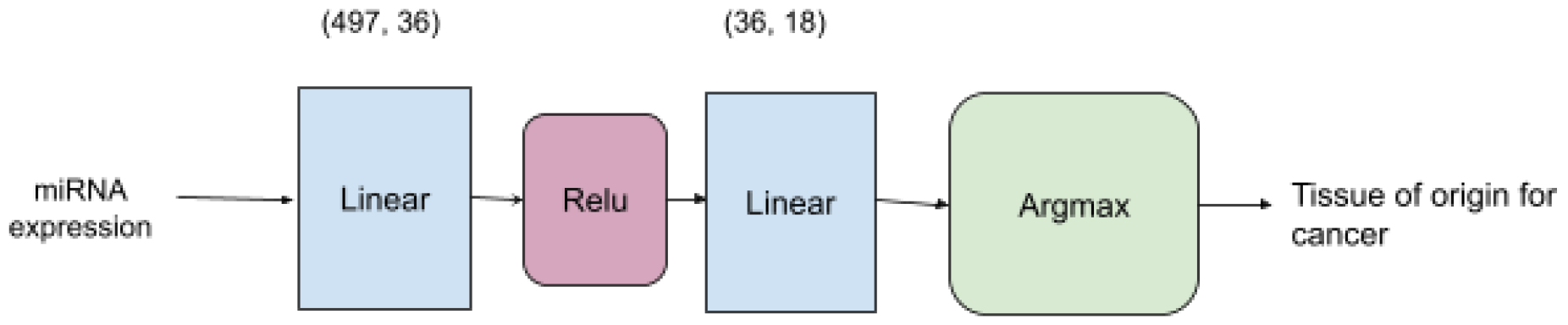
Deep Learning model architecture. A schematic of the machine learning model architecture.

We used dropout for the input layer [22] as it is a common technique to improve model accuracy and reduce overfitting. We also augmented our input data with noise.

To evaluate the performance of our models, we computed confusion matrices for performance on metastatic samples (Figure 2A,B) and plotted the ROC curves for performance on metastatic skin cancer (Figure 2C,D), as the majority of the metastatic samples were obtained from skin cancer cases. We observed that the deep learning model performed significantly better than our decision tree model, which was consistent when evaluated on the SRA validation cohort (Figure 3).

**Figure 2:**
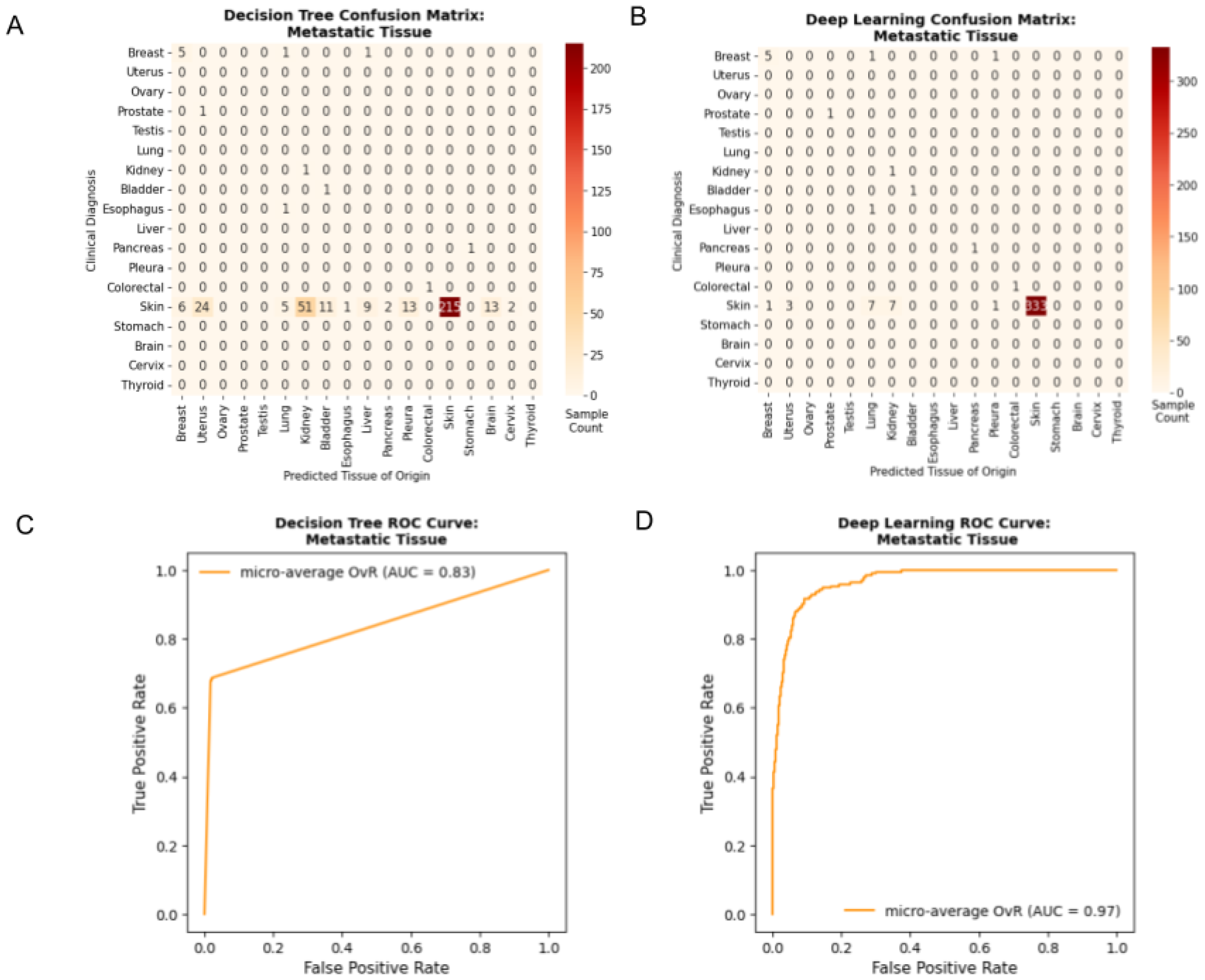
Representative Confusion Matrices and ROC Curves for Created Models on Metastatic Samples. A) Confusion Matrices for Decision Tree and B) Deep Learning model on metastatic samples. C) ROC curves for Decision Tree and D) Deep Learning model on metastatic samples.

**Figure 3:**
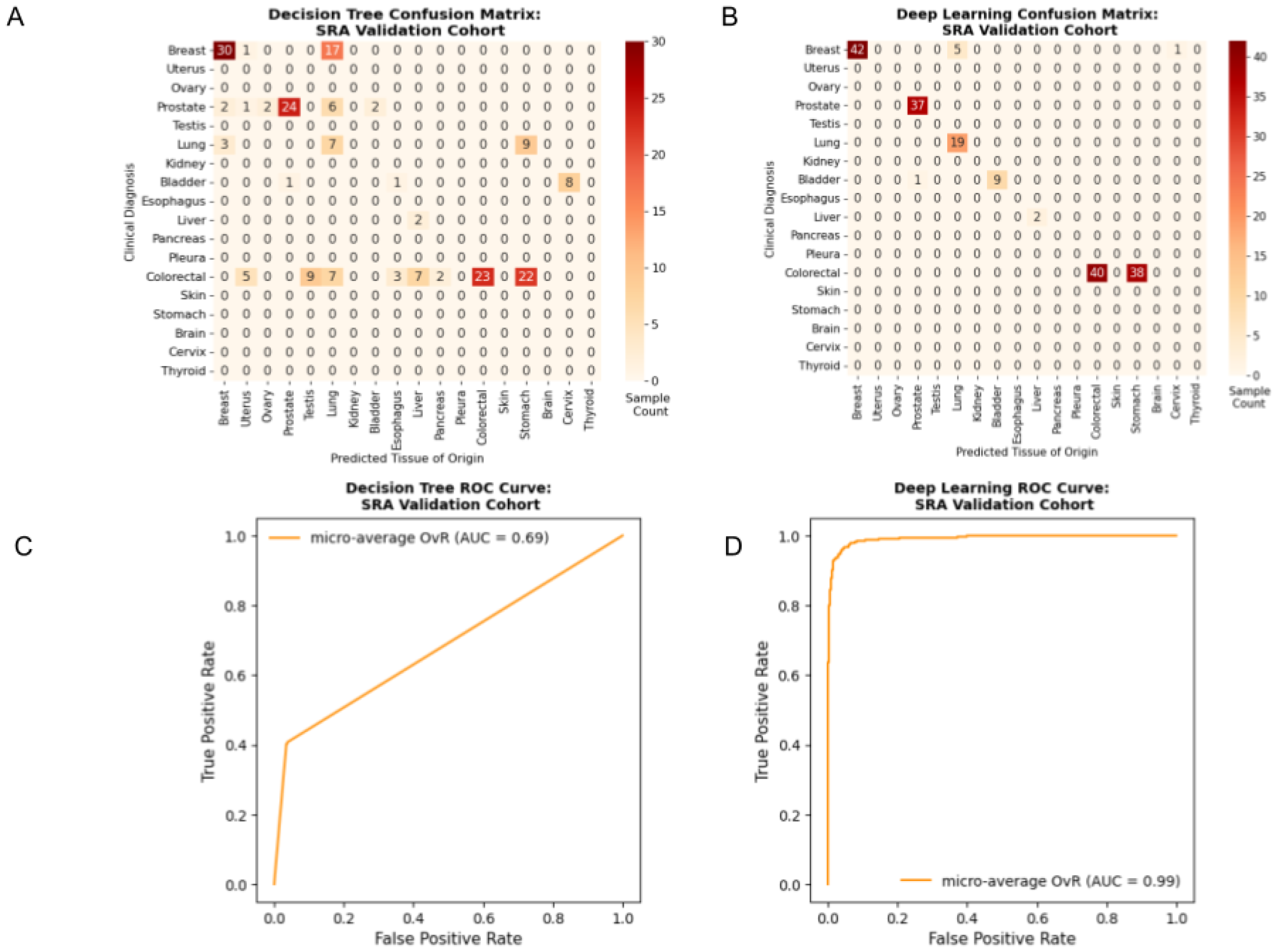
Representative Confusion Matrices and ROC Curves for Created Models on SRA Samples. A) Confusion Matrices for Decision Tree and B) Deep Learning model on samples from SRA. C) ROC curves for Decision Tree and D) Deep Learning model on samples from SRA.

These results confirm our hypotheses and show that we were able to predict the tissue of origin with high accuracy using deep learning. Furthermore, our findings demonstrated that deep learning techniques significantly increase the accuracy in comparison to decision tree, logistic regression and random forest models.

To reveal the significance of individual features, we performed feature importance analysis using permutation feature importance method (Figure 4). The top three microRNAs contributing to our deep learning model based on our combined normal and primary site training set are mir-10b, mir-196, and mir-205. Mir-10b has been shown to function as a metastasis promoting factor in many cancer types. In fact, it was one of the first microRNAs to have been discovered with aberrant expression in cancer cells [8]. Mir-196 has been linked to the progression of many cancers, notably metastatic colorectal cancer [9] while mir-205 expression is downregulated in metastatic breast and prostate cancer [10].

**Figure 4:**
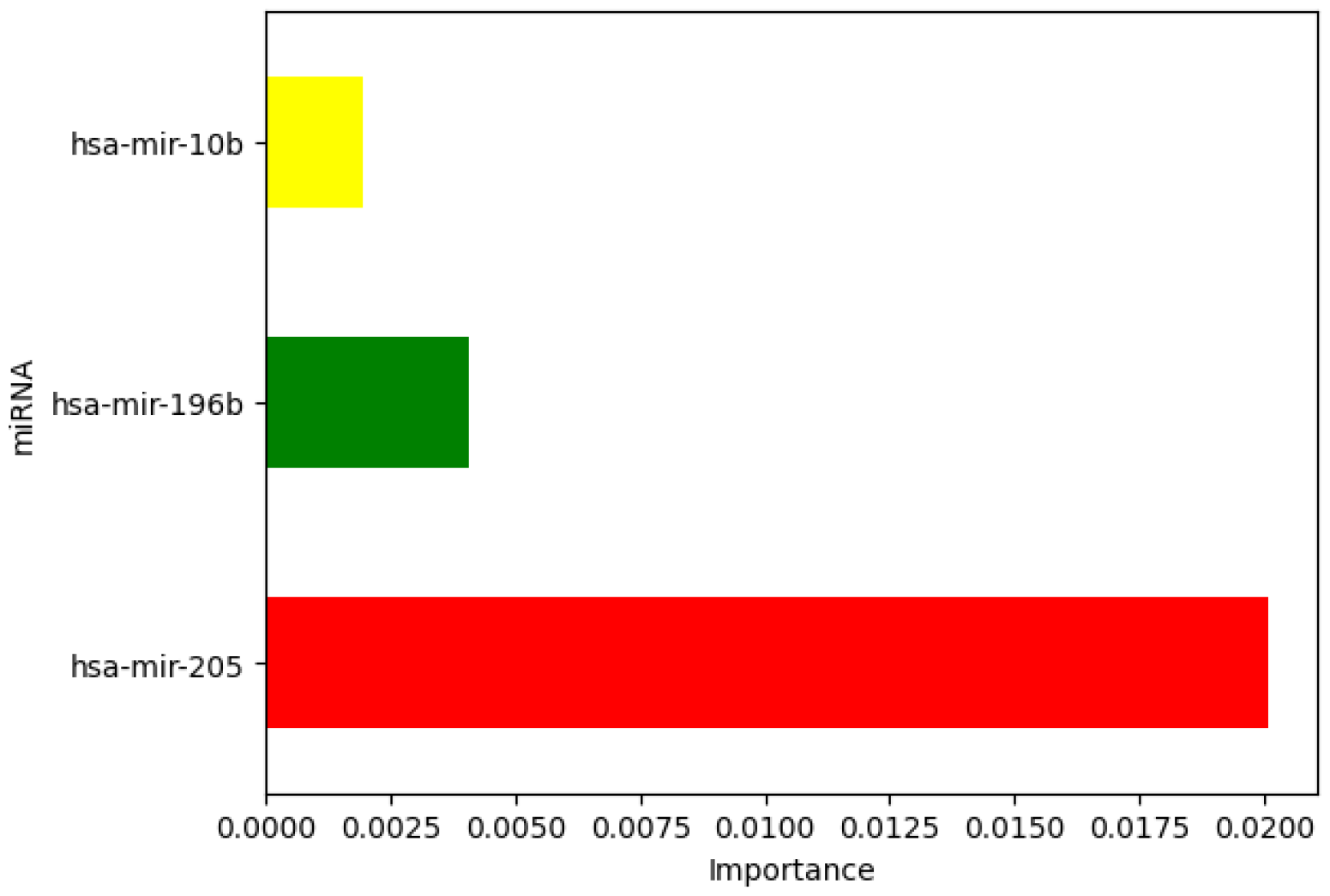
Permutation feature importance for the top three microRNA candidates. A bar graph of the importance values for the three top microRNA candidates in the Random Forest model.

In Figure 5A, we see that there is a clear distinction in miRNA expression based on the type of cancer. For example, in the row corresponding to hsa-mir-196b, we see clear variation in the level of expression for different types of cancer, most notably colorectal and thyroid. Figures 5B and 5C show the PCA and t-SNE visualization of data corresponding to the six cancer types with the most samples in our dataset, using only the top 10 miRNA features. In the PCA plot, note that there is significant overlap between the cancer types, while in the t-SNE plot, the cancer types are well separated. This suggests that even with just ten miRNA features, machine learning models will be able to correctly grasp patterns and predict tissue of origin.

**Figure 5:**
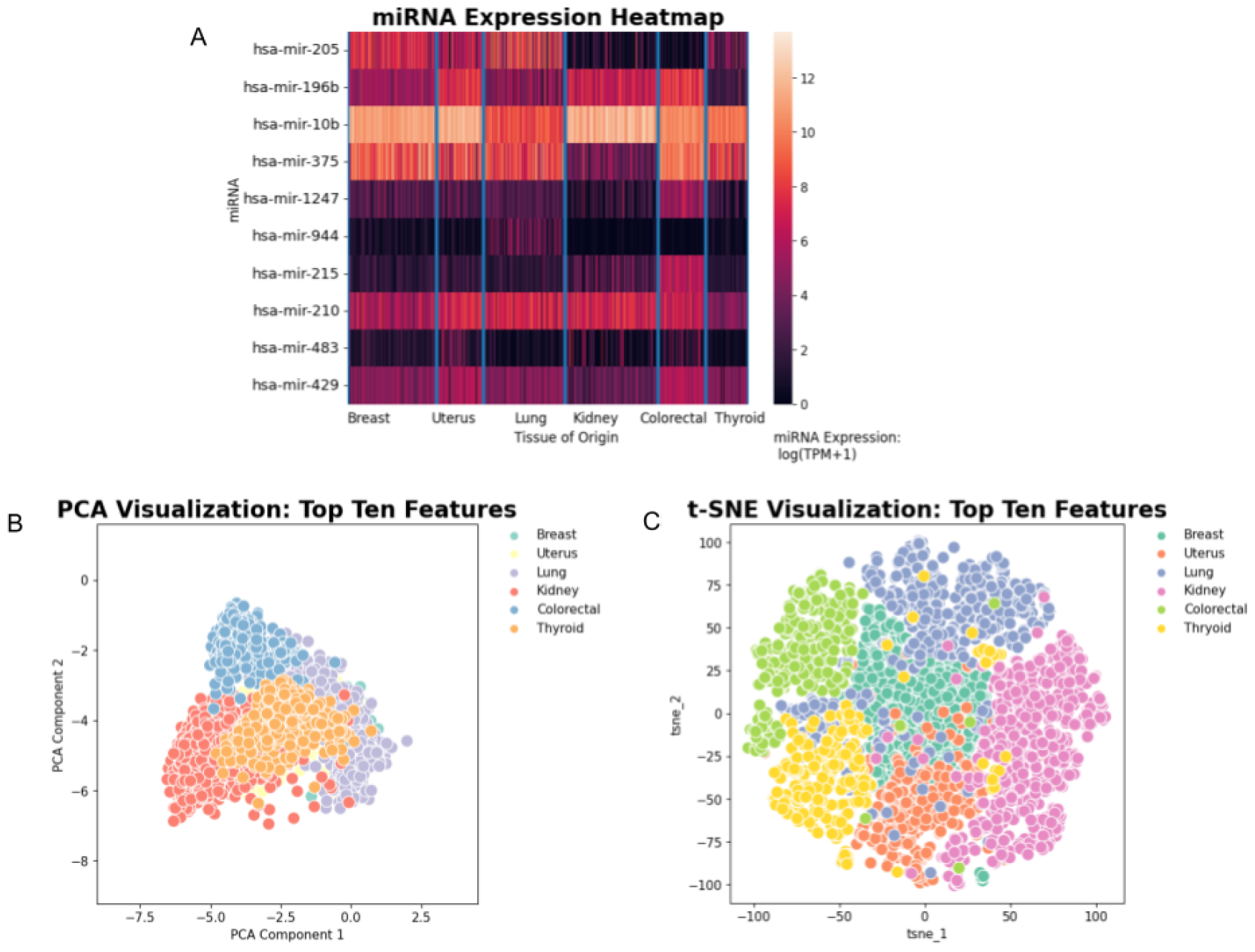
miRNA Expression Heatmap (A), PCA Visualization (B), TSNE Visualization (C)

## DISCUSSION

In these investigations while employing successively more powerful classifiers, we were able to detect the tissue of origin on solely metastatic cancer samples with accuracies ranging from 62.5% with a decision tree to 96.1% with a deep learning model. Results show that models trained only on primary tumor and solid tissue normal data are able to predict tissue of origin from metastatic gene expression data. Our methods have also identified promising biomarker candidates, reaffirming prior research in this field and demonstrating the potential of machine learning. In order to verify the robustness of our model, we assessed its performance on primary tumor data from the SRA and obtained accuracies ranging from 41.2% with decision tree to 81.2% when employing deep learning.

In the deep learning ROC curves (Figure 2C,D and Figure 3C,D), we see that the true positive rate is close to 1 when the false positive rate is close to 0. This signifies that our model exhibits a high level of accuracy and a minimal error rate. In this context, errors in classification come at a significant cost. Errors in detecting tissue of origin can lead to poor treatment outcomes if an ineffective therapy is chosen based on the results. When comparing the confusion matrices for the decision tree and deep learning model on the metastatic test set, we can see very significant improvement. Particularly, in the deep learning model, the confusion matrix is nearly diagonal, implying that our model is correctly classifying tissue of origin in most instances. The cancer types with the most notable improvement compared to the decision tree model (Figure 2A,B) were metastatic skin cancer.

The predominant failure of our model on our validation SRA cohort was within colorectal cancer as can be seen in Figure 2. Many colorectal samples were incorrectly classified as stomach/gastric cancer. This is interesting to see as colorectal and stomach cancer are often synchronous with probabilities ranging from 20.1% - 37.2% [23].

We used permutation feature importance, a model agnostic metric that permutes features across samples in the test set to assess the change in model accuracy. The results are in line with existing research in this area and serve as a good indicator of the feasibility of machine learning techniques to identify promising biomarkers.

### Limitations

To effectively utilize our model in clinical care, accuracy must be improved further. Our model currently performs with an accuracy of 96.1%. While this may seem impressive, clinical classifiers should be highly accurate so that there are a negligible number of cases with errors in identifying tissue of origin. To improve the accuracy, we will need much larger datasets, and as the non-coding genome continues to reveal significant contributions to cancer, we predict that available datasets will expand. A further limitation to our study was that the available metastatic datasets are predominantly skin cancer. Thus, access to a larger, more varied, dataset would improve our assessment of model performance. Furthermore, in order to develop a truly noninvasive method of tissue of origin identification relevant for all cancers, it would be ideal to extend our method to microRNA expression data from blood samples. Detecting the tissue of origin through blood-based microRNA biomarkers would significantly impact the diagnosis and treatment of CUP patients.

## Conclusions

To summarize, our developed machine learning models can accurately identify the tissue of origin with high accuracy from microRNA expression data when trained on primary tumor and solid tissue samples. Importantly, our results identified key microRNA biomarkers of tissue type. Our models are robust and perform well across different datasets (TCGA and the SRA dataset). We look forward to developing further deep learning models that can accurately detect tissue of origin as microRNA datasets expand, with the goal of having a non-invasive test for diagnosing the presence of cancer and determining the cancer tissue of origin with high accuracy.

## ACKNOWLEDGMENTS

We are grateful to the Cancer Genome Atlas Project (TCGA) and patients for providing the data used in this research. We are thankful to Soroush Hajizadeh for reviewing our code and providing insightful feedback. The results are in part based upon data generated by the TCGA Research Network [6].

## SUPPLEMENTAL MATERIALS

Figure S1: PCA and TSNE analysis for all miRNA features.

Figure S2: Schematic of data preprocessing pipeline.

Table S1: Distribution of tissue of origin across datasets.

**Figure S1:**
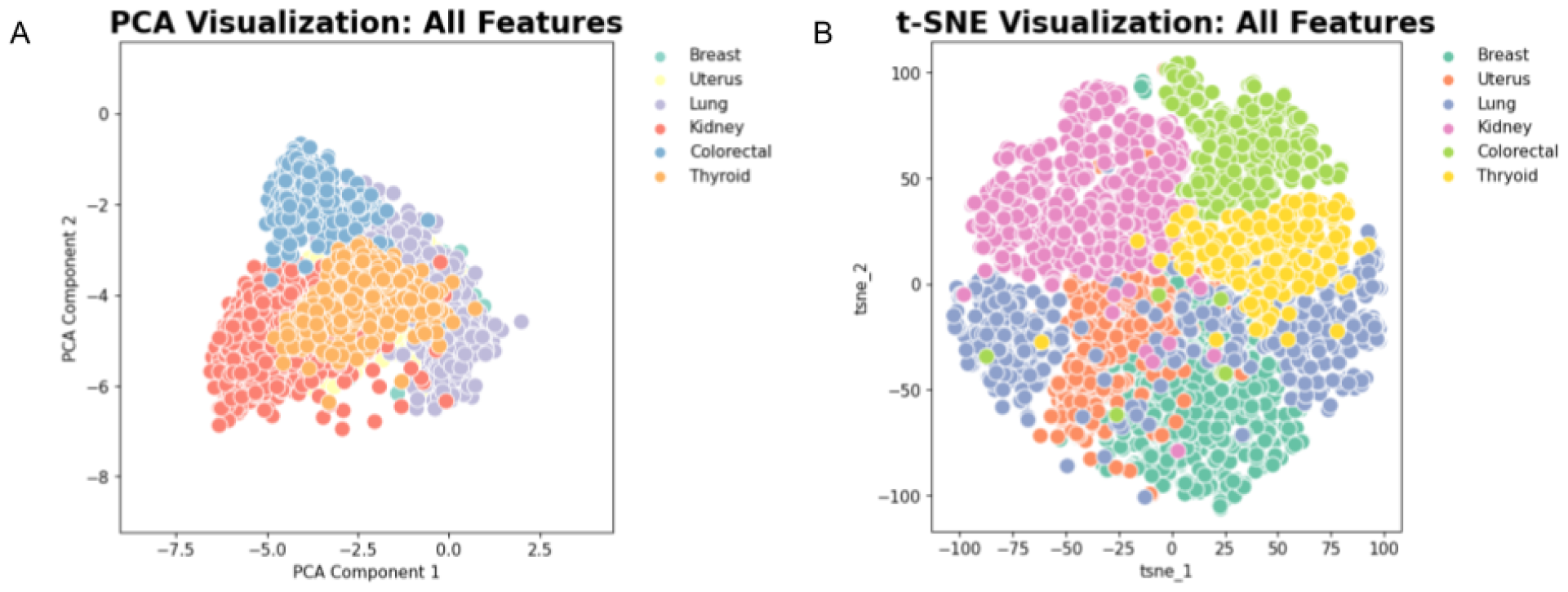
Figure S1A and S1B show the *PCA and t-SNE visualization of data corresponding to six cancer types, using* ***all*** *miRNA features. These plots are comparable to those shown in Figure 5B and Figure 5C, showing that the top 10 miRNA features contain most of the relevant information. This shows that the permutation feature importance has successfully identified a subset of the most relevant miRNA features*.

**Figure S2:**
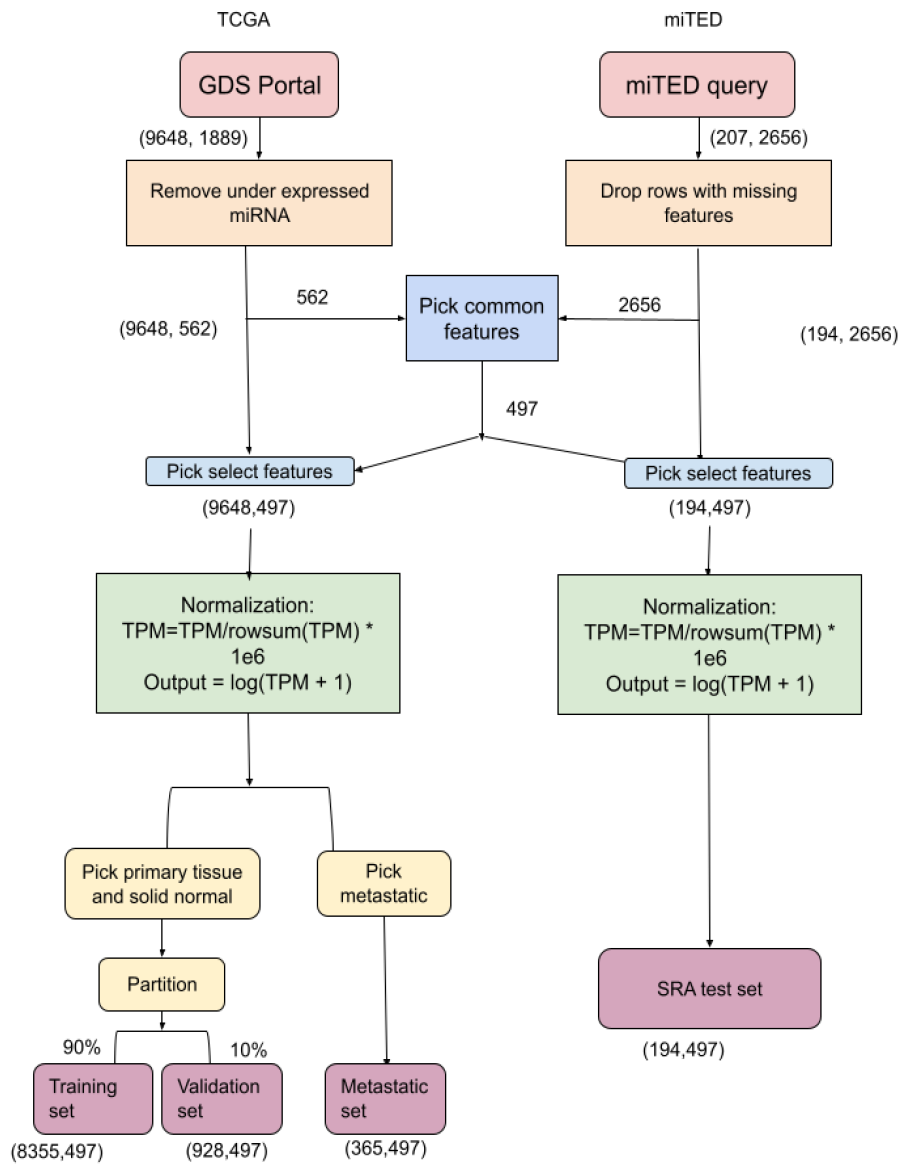
Overview of our data processing pipeline. We first obtained data from the GDS portal and miTED website. We removed under expressed microRNA from the TCGA (GDS) data and removed samples containing missing features from the miTED data. Common features were selected between both datasets, reducing the number of microRNA to 497. On both datasets, we first normalized the TPM of the selected features per sample to sum to a million. We then performed a log transformation to restrict the range of microRNA expression to the model. The TCGA dataset was split into the primary tissue and solid normal combined set and the metastatic test set. The combined set was further split into a training and validation set.

**Table S1:** Distribution of cancer types in dataset

| Tissue of origin | Primary | Solid | Metastatic | SRA |
| --- | --- | --- | --- | --- |

**Table S1:** *Distribution of cancer types in dataset*
|  | Tumor | Tissue |  |  |
| --- | --- | --- | --- | --- |
| Breast | 1096 | 104 | 7 | 48 |
| Uterus | 602 | 33 | 0 | 0 |
| Ovary | 490 | 0 | 0 | 0 |
| Prostrate | 498 | 52 | 1 | 37 |
| Testis | 150 | 0 | 0 | 0 |
| Lung | 997 | 91 | 0 | 19 |
| Kidney | 1093 | 142 | 1 | 0 |
| Bladder | 417 | 19 | 1 | 10 |
| Esophagus | 186 | 13 | 1 | 0 |
| Liver | 372 | 50 | 0 | 2 |
| Pancrea | 178 | 4 | 1 | 0 |
| Pleura | 87 | 0 | 0 | 0 |
| Colorectal | 616 | 11 | 1 | 78 |
| Skin | 97 | 2 | 352 | 0 |
| Stomach | 446 | 45 | 0 | 0 |
| Brain | 512 | 5 | 0 | 0 |
| Cervix | 307 | 3 | 0 | 0 |
| Thyroid | 506 | 59 | 0 | 0 |

## Notes

https://www.cancer.gov/ccg/research/genome-sequencing/tcga

https://dianalab.e-ce.uth.gr/mited/#/

https://github.com/Anisha234/Identifying-Tissue-of-Origin-From-miRNA

### Competing Interest Statement

The authors have declared no competing interest.

